# Effect of Schizophrenia linked *DRD2* gene mutations and correlation of social behavior with biochemical changes in a post-weaning isolated mouse model

**DOI:** 10.1101/2023.11.02.565337

**Authors:** Nivedhitha Mukesh, Pavan Kumar Divi, Shimantika Maikap, Saswati Mohapatra, Muskaan Rajan, Sanjana Kanthimath, Nishant Mishra, Karunakar Kar, Bibin G Anand, Anil Annamneedi

**Affiliations:** Department of Biotechnology, School of Bioengineering, SRM Institute of Science and Technology, Kattankulathur-603203, Tamil Nadu, India; Biophysical and Biomaterials Research Laboratory, School of Life Sciences, Jawaharlal Nehru University, New Delhi-110067, India

**Keywords:** Neurological disorders, ASD, Schizophrenia, Epilepsy, Indian Population, Post-weaning social isolation mouse model

## Abstract

Neurological disorders encompass a diverse range of conditions that affect individuals cognitive, emotional, and social functioning. Though these disorders are multifactorial, genetic factors play a significant role in the pathogenesis. Especially, synaptic gene mutations or synaptic protein dysfunction are shown to be closely associated with neuropathology. This study aims at understanding the critical role of synaptic compartment by conducting comprehensive analysis of existing synaptic gene mutations responsible for the development of three significant disorders in the Indian population: autism spectrum disorder (ASD), epilepsy and schizophrenia (SZ). Our *in-silico* analysis predicts that mutations in synaptic genes *RPL10* (rs387906727), *GABRA1* (rs121434579) and *DRD2* (rs1801028) corresponding to ASD, epilepsy and SZ respectively, are deleterious. Of these, SZ-related mutations in *DRD2* (D(2) dopamine receptor) are deleterious and due to its genetic association also with ASD as well sociosexual behavior, this study focuses on DRD2. *In silico* analysis using molecular docking revealed an abnormal interaction between D(2) dopamine receptor and neuronal calcium sensor 1, which may hamper neurotransmitter regulation. We further employed a post-weaning social isolation mice model of SZ to investigate the D(2) dopamine receptor expression levels in hippocampus and social behavioral changes. We observed a reduced immunofluorescent intensities of D(2) dopamine receptor and NCS1 compared to group-housed controls in hippocampus and a trend towards an impaired sociosexual behavior characterized by anogenital sniffing in a male-female social interaction test. Altogether, our study helps to further our understanding of synaptic signaling in the context of SZ and decode the probable mechanism by which disrupted synaptic signaling and protein-protein interaction may lead to the disease pathology and further aid in identifying novel therapeutic targets.

## 1 Introduction

The brain carries out several signal transduction pathways critical in controlling all voluntary and involuntary actions, cognition, perception, judgement, memory, and retrieval among other diverse biological functions. Synaptic communication is critical for the transfer of information and the neurological disorders that manifest due to synaptic dysfunction are broadly termed synaptopathies (1). Genetic mutations in genes coding for proteins ultimately change protein structure, function, interaction with protein partners and may contribute to a phenotypic attribute of the disorder. As a result, studying synapses at the molecular level is critical in determining etiology.

Prevalence of various neurological disorders including autism spectrum disorder (ASD), epilepsy, schizophrenia (SZ) etc. is increasing in India during the last decades (2–4). ASD and SZ are two different neurological conditions, with distinct phenotypic changes, yet both display disturbed social cognition, altered neuroanatomy, disturbed GABAergic functioning, and co-occurrence of epilepsy (5–7). Human mutations in several of the genes encoding for synaptic proteins have been identified across all these conditions and potentially involved in disturbance of convergent pathways at synaptic compartments. Understanding the shared origins and fundamental distinctions in disorders linked with synaptic dysfunction could lead to novel therapeutic approaches.

Post-weaning social isolation (SI) in mice is an established and widely employed mouse model to study changes related to development of neuropsychiatric conditions including ASD and SZ (8). Previous studies employing the SI model mice display long-lasting changes in brains of isolated mice including SZ-related behavior, altered functioning of dopaminergic, serotonergic, glutamatergic systems and morphological changes in different brain areas, strongly resembling the SZ core features (8,9).

Our current study focuses on analyzing the existing gene mutations linked to ASD, epilepsy and SZ with a special focus on the Indian population, their effect on protein structure, and protein-protein interaction through *in silico* analyses. Further, using the SI model of SZ, we aim to understand the biochemical alterations in hippocampus and correlation with the social behavior.

## 2 Materials and methods

### 2.1 *In silico* analysis

Primary list of all mutated genes associated with neurological disorders such as ASD, epilepsy and SZ in the Indian population was gathered from PubMed and Google Scholar (Supplementary Table 1). Using the SynGo database (10) the list of genes was segregated into synaptic and nonsynaptic genes, and only synaptic genes were considered for further analysis. The SNPs in synaptic genes were further sorted into exonic (synonymous and non-synonymous) and intronic based on position of occurrence. To sift out highly impactful mutations, Position-Specific Independent Counts Scores (PSIC) tools such as PolyPhen2 and SIFT were used (11). SIFT (Sorting Intolerant From Tolerant) uses sequence homology to predict whether an amino acid substitution will affect protein function or not. PolyPhen scores can be interpreted as follows: 0.0 to 0.15 - predicted to be benign; 0.15 to 1.0 - possibly damaging. 0.85 to 1.0 - confidently predicted to be damaging. SIFT score uses the same range, 0.0 to 1.0, but in reverse. We next compared all the synaptic genes to identify the most deleterious gene mutation per neurological condition. The nucleotide sequence of the respective gene variants that are disease-causing were taken from NCBI (the gene code for *RPL10* is 6314, for *GABRA1* is 2554 and for *DRD2* is 1813). The mutations were induced by altering the nucleotide sequence (SNP) according to the respective mutations. Amino acid sequences were identified for the wild type and mutated gene-encoded protein using Mutation Taster (12). The 3-D protein structures of the wild-type and mutant variants were generated using SWISS-MODEL (13). Molecular dynamics (MD) simulations were conducted using GROMACS version 2023.1 (14) with ff19SB (15) Amber force field and TIP3P water model. The proteins were centered in a 10nm simulation box, solvated, and neutralized by counter ions. First, proteins underwent minimization, followed by 2 staged equilibrations—NVT and NPT equilibration employing V-rescale (16) thermostat and Parrinello-Rahman (17) pressure coupling algorithms. The final 100ns production was run at physiological temperature (310K), from which frames at 0ns and 100ns were extracted for subsequent analysis using GROMACS built-in tools. Furthermore, snapshots and interactions were visualized and analyzed using Discovery Studio Visualizer (BIOVIA, 2021).

The list of interacting protein partners to the wild type (WT) protein were prepared using the STRING (Search Tool for the Retrieval of Interacting Genes/Proteins) analysis tool (18), with highest confidence (0.900) as minimum required interaction score. To check for the binding differences between the wild type (WT) protein and mutated (MT) proteins with their interaction partners ClusPro (19) was used. Molecular docking was done to study the interacting pair with the largest change in binding energy to study the alteration in protein-protein interaction.

### 2.2 Animals

4-week-old weaned male C57BL6/J mice were obtained from a commercial breeder (MASS Biotech, Chennai, India) and subjected to different housing conditions. The social isolated (SI) group of mice were raised in an individual cages for 4 weeks and compared with the group housed (5 per cage) control mice (20). Behavioral testing and biochemical analyses were carried out at an age of 9 weeks old.

All mice were housed, and behavior analyses were carried out, in Class 100000 clean room environment at our animal house, SRM-DBT platform, SRM Institute of Science and Technology. Mice were provided with food and water *ad libitum*. Temperature was maintained at 22 ± 1°C, relative humidity was maintained at 50 ± 5 % 3 and 12 hour Light:Dark cycle wad followed. The experimental protocol was approved by the Institutional Animal Ethics Committee.

### 2.3 Social behavior

Male-female interactions were carried out in an open field arena. Female C57BL6/J mice at an age of 8 weeks were used. The test male mice were habituated to the context prior to carrying out the behavioral assay for 5 minutes. Post the female mouse’s introduction and habituation to the arena for 2 minutes, the male test mouse was introduced, and social interactions (anogenital sniffing) were recorded for 3 minutes. Protocol was modified from previous study (21). Experiment was conducted on two cohorts of mice. Cohort 1 (5 group housed and 5 SI mice) does not receive any additional stressor, whereas cohort 2 (5 group housed and 5 SI mice) received a mild stressor-intraperitoneal (I.P.) injection of DMSO (1.5%) 30 min prior to behavioral testing (22). Parameters like following behavior and anogenital sniffing were analyzed using the BORIS (23) software.

### 2.4 Immunohistochemistry (IHC)

Brains were isolated post cervical dislocation of overanastetized mice and fixed with 4% PFA overnight. Brains were thsen cryoprotected in sucrose solution and stored at −20 °C until used for immunohistochemical staining, which was performed as described previously on 30 μm sagittal brain sections (24). Antibodies used includes Rabbit anti-DRD2 (1:200, ABclonal, Catalogue #A12930; RRID:AB_2759776), Rabbit anti-NCS1 (1:200, ABclonal, Catalogue #A13568; AB_2760448) and Goat anti-Rabbit IgG Alexa Fluor 488 (1:200, ABclonal, Catalogue Number #AS053; RRID:AB_2768320). Fluorescent images were obtained employing Leica DM6 Fluorescent Microscope using 10X objective and the integrated density values were measured in hippocampus using the ImageJ (NIH, http://rsb.info.nih.gov/ij/).

### 2.5 Statistical analysis

GraphPad Prism (GraphPad Software Inc., version 9) was employed for statistical analysis of behavior and quantitative IHC. Normality test (Shapiro-Wilk test) was performed on behavioral data before commencing statistical tests. Paired or unpaired Students *t* tests were used. In case the data did not pass the normality test, Mann-Whitney *U* test was used. Unpaired parametric student t-test was carried out to analyze the IHC data. Data are represented as means ± standard errors of the mean (SEM), with probability values of p < 0.05 were considered as statistically significant.

## 3 Results

### 3.1 Synaptic gene mutations implicated in the Indian population

Using the SynGO (cellular component) database, we narrowed down to synaptic genes from the list of genes whose mutations are implicated in the Indian population with neurological disorders. Indeed, except for Epilepsy conditions, the synaptic genes are localized to both pre and post synaptic compartments in ASD and Schizophrenia (Figure 1 and Supplementary Table 1). Intronic variants, despite not contributing to changes in the coding sequence, have been implicated in various disorders (25). Surprisingly, more than 50% of the mutations implicated in Schizophrenia, both synaptic and non-synaptic genes are intronic variants, whereas we did not observe this trend in case of ASD and Epilepsy (Supplementary Table 1).

**Figure 1:**
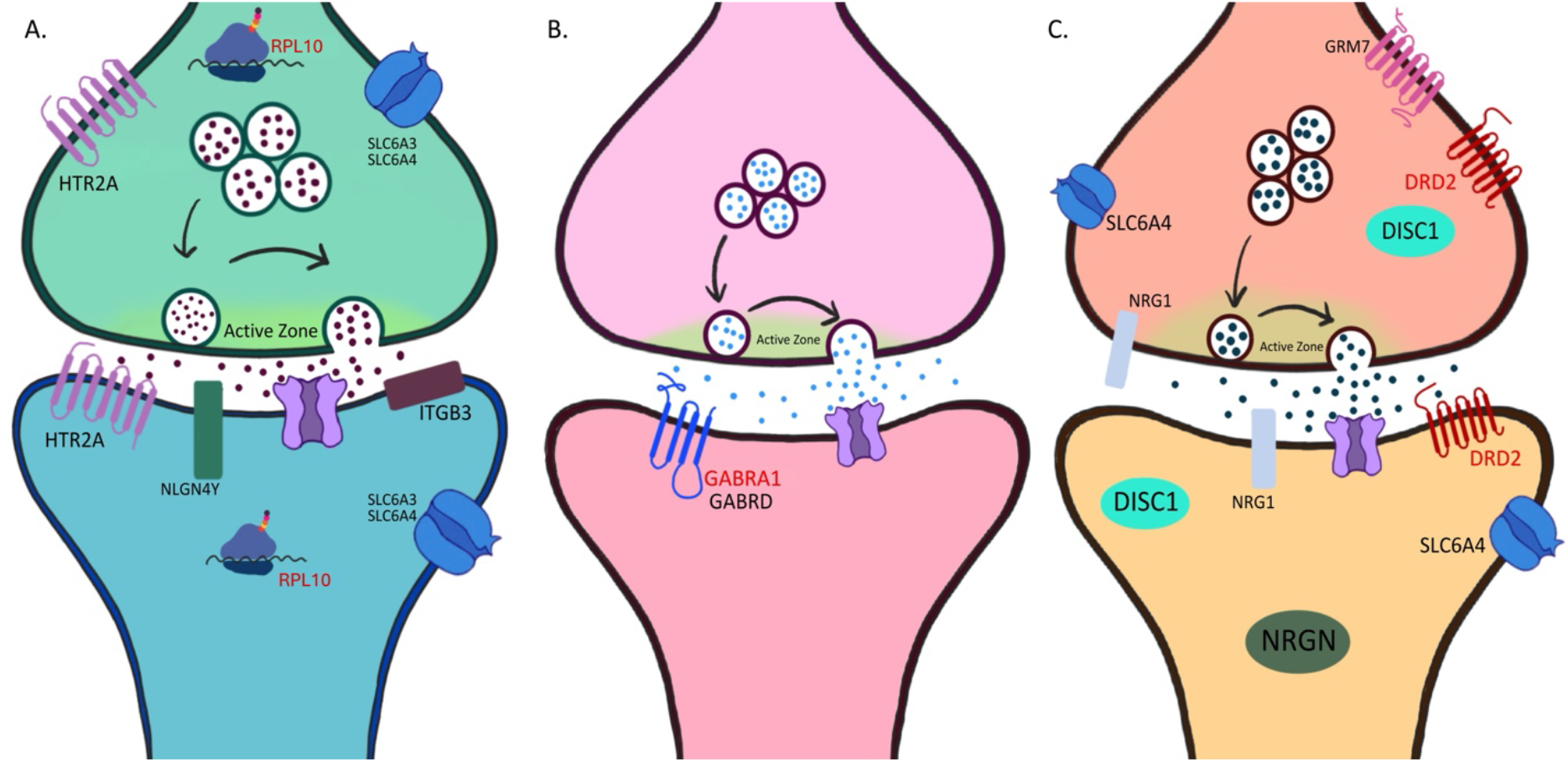
Localization of synaptic genes. The list of genes identified from various studies conducted on the Indian population with respect to neurological conditions such as ASD **(A)**, epilepsy **(B)** and schizophrenia **(C)**, are stratified into synaptic genes, both pre and post synapse using SynGO-cellular component. Most affected gene is highlighted in red.

### 3.2 Severity of impact of mutations on protein structure

First, we identified potential genes that are most deleterious in each condition, using bioinformatic prediction tools. We concluded that the three genes *RPL10, GABRA1, DRD2* and their mutations associated with ASD, epilepsy and SZ, respectively, are of highest impact potential (Table 1). All the three genes are predicted to be damaging or deleterious based on the PolyPhen-2 and SIFT scores. Among the three genes (where *RPL10* encodes for Large ribosomal subunit protein uL16, *GABRA1* encodes Gamma-aminobutyric acid receptor subunit alpha-1, *DRD2* codes for and D(2) dopamine receptor). *GABRA1* and *DRD2* encoded proteins are vital in neurotransmission related signal transduction pathways and are highly impaired in the diseases they are associated with, whereas *RPL10* codes for a ribosomal protein critical for the assembly of the polysomes (26–28).

**Table 1:**
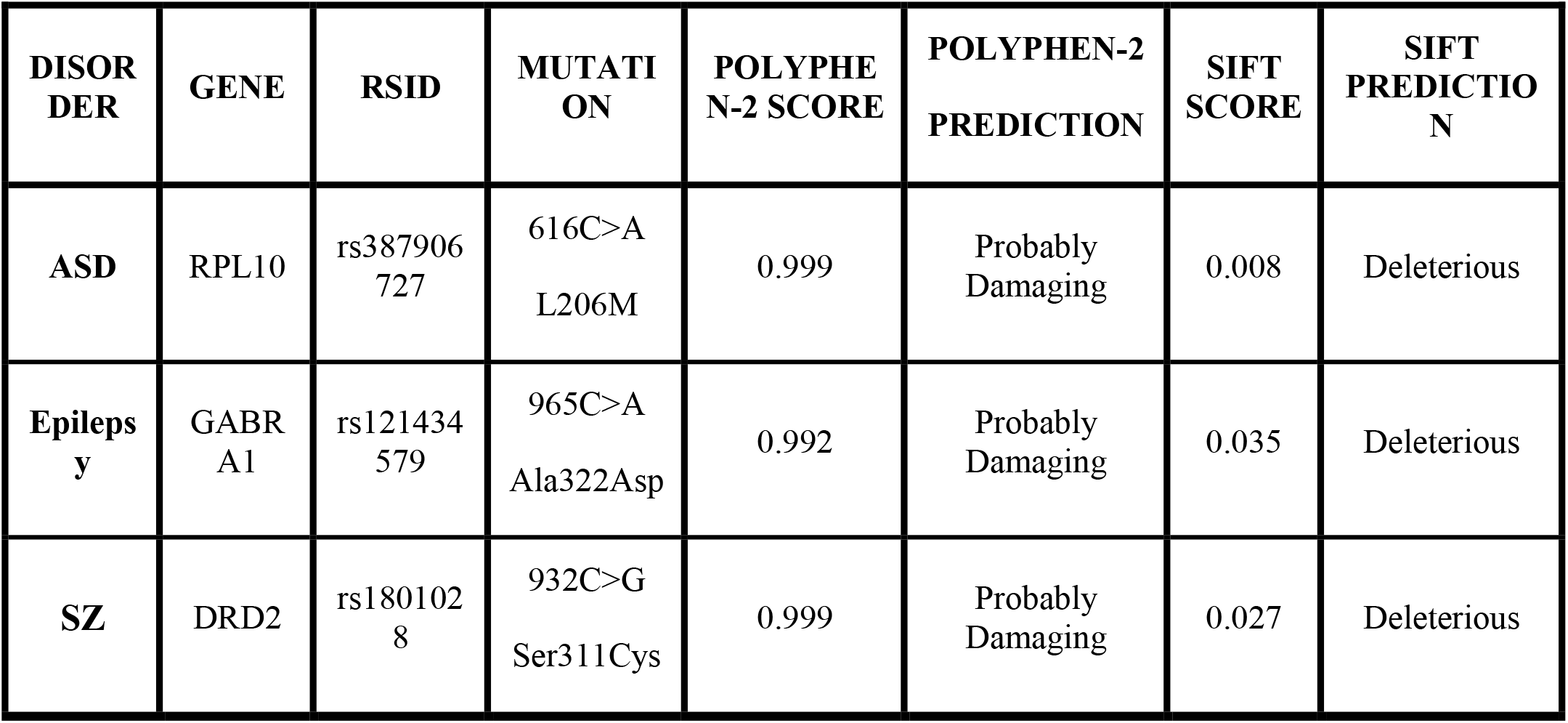
Position-Specific Independent Counts Scores. Determination of severity of the mutation and its impact on the structure and function of respective protein using the PSIC tools Polyphen-2 and SIFT.

### 3.3 Molecular Dynamics Simulation

Further, we sought to visualize the structural differences in proteins coded by mutated genes in comparison with WT (Supplementary Figure 1). The sequence identity for the aforementioned proteins are as follows: Large ribosomal subunit protein uL16 - 98.9%, Gamma-aminobutyric acid receptor subunit alpha-1 - 99.07%, and D(2) dopamine receptor - 99.77%.

Post analysis of our simulation studies revealed that the DRD2 WT model, in the 0ns frame the Ser311 residue falls in the disallowed region but falls in the beta turn conformation in the 100ns frame (Figure 2A). In the MT model, the Cys311 residue falls in the coil conformation region at 0ns but shifts to β-turn region in the 100ns frame (Figure 2B). However, we did not find any significant alterations in protein structure due to mutations in either RPL10 or GABRA1 and we further focused on analyzing the structural changes in DRD2.

**Figure 2:**
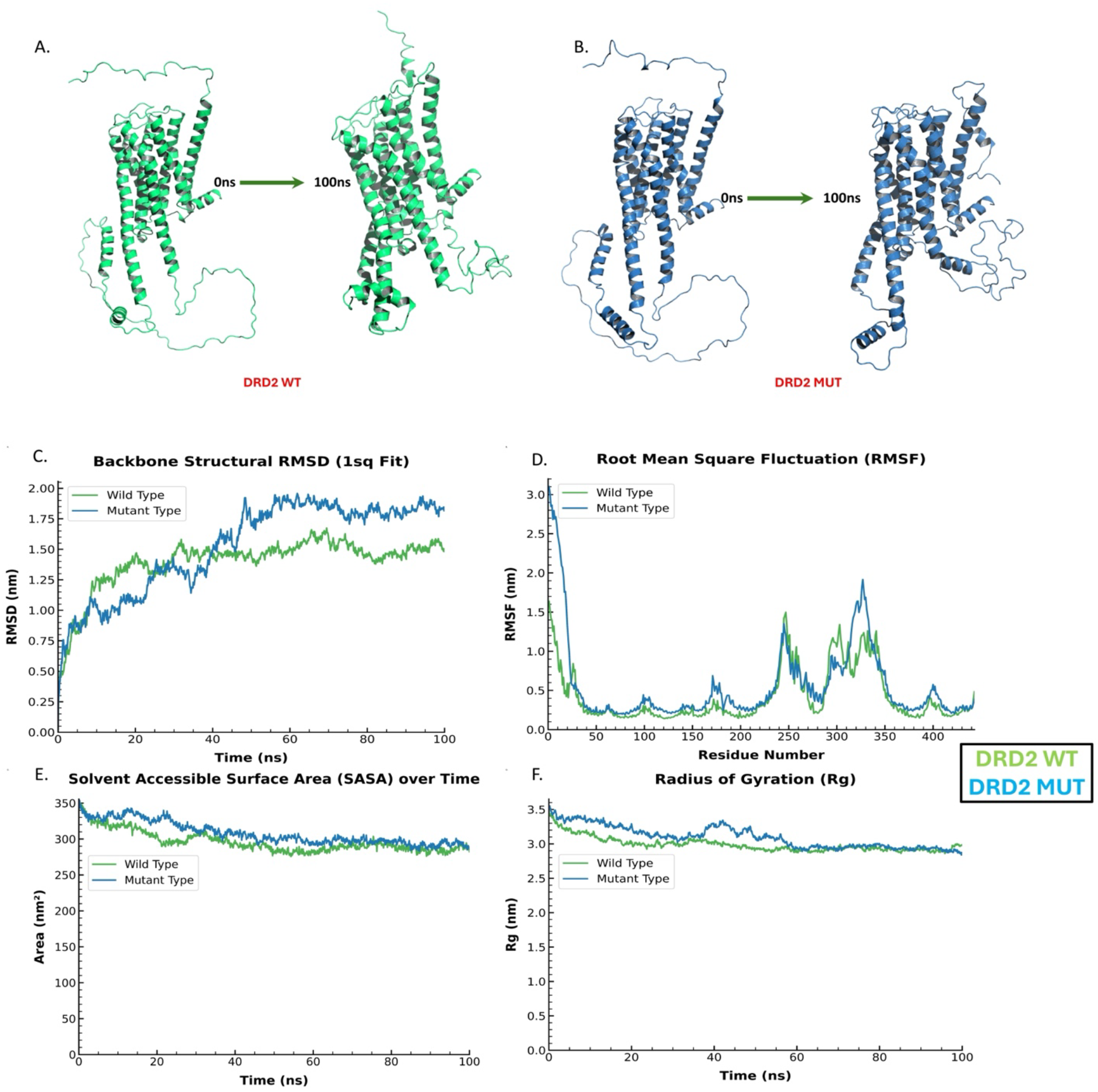
Molecular Simulation Dynamics of Dopamine Receptor D2. The figure shows the 0th nanosecond and 100th nanosecond snapshot in the molecular dynamics simulation of WT **(A)** and MT **(B)** Dopamine Receptor D2. **(C)** Root mean square deviation vs time graph. **(D)** Root mean square fluctuation of amino acids vs residue number graph. **(E)** Solvent accessible surface of protein molecule vs time graph **(F)** Radius gyrates of simulated system vs time graph.

Despite the secondary structure both being β-turn in DRD2, the phi angle is observed to change significantly. This shows that there are significant structural constraints due to the amino acid substitution. In the root mean square deviation (RMSD) graph for DRD2 (Figure 2C) after the 45ns of simulation, there is a higher degree of fluctuation in the WT protein model than the MT protein model, indicating a higher rigidity in MT than the WT model. In the root mean square fluctuation (RMSF) graph of DRD2 (Figure 2D), the Ser311 residue in the WT model falls in a higher peak region of the RMSF graph than Cys311 in the MT model, indicating relatively higher-level fluctuation in its position in the backbone in comparison with the MT model, thus supporting the hypothesis that the MT model is relatively more rigid than the WT model. Such phenomena may be due to the residues interacting spatially with free cysteines of non-membrane proteins, revealing a predominant association with hydrophobic residues and a lesser affinity for polar or charged residues, thus being less flexible/dynamic. The solvent-accessible surface (Figure 2E) and radius of gyration (Figure 2F) for both the models is similar within normal ranges of fluctuation.

### 3.4 Impact of mutation on protein-protein interaction

It is well understood that point mutations significantly contribute to stabilizing protein structures, as demonstrated in studies on lysozyme (29). Therefore, we anticipate that similar mutational alterations in our protein may also impact its stability. In this study we have identified the interaction partners of each of the impacted proteins in WT condition (Figure 3 A-C), using the STRING tool. From this list, we have assessed the most affected “interacting partner-POI (protein of interest) interaction” (Table 2). In the case of ASD, the interaction between RPL18A and the Large ribosomal subunit protein RPL10 is highly affected (difference in weighted scores between WT and MT conditions is -48.6), which might result in disruption of polysome formation. For epilepsy, the affected interaction is between GABBR2 and GABRA1 (difference in weighted scores is -375.7). Finally, we identified that the strongly affected interaction is between NCS1 and D(2) dopamine receptors, which is implicated in SZ, difference in weighted scores is 890.1 (Table 2).

**Table 2:** Protein-protein interaction scores (Lowest Energy). Analysis of the impact of mutation on protein-protein interaction, using ClusPro and the change in lowest energy scores. Most altered score is highlighted.

| Disease | Protein of Concern | Interacting Protein (Gene) | Weighted Score-Wild Type Protein | Weighted Score - Mutated Protein | Difference |
| --- | --- | --- | --- | --- | --- |
| ASD | Large ribosomal subunit protein uL16 (rs782521991) | *RPL13* | -887.8 | -888.4 | 0.6 |
|  |  | *RPL15* | -765.6 | -765.9 | 0.3 |
|  |  | *RPL18A* | -844.7 | -789.3 | -55.4 |
|  |  | *RPL19* | -727.2 | -680.1 | -47.1 |
|  |  | *RPL35* | -680.4 | -654 | -26.4 |
|  |  | *RPL8* | -904.2 | -856.5 | -47.7 |
|  |  | *RPS11* | -741.4 | -741 | -0.4 |
|  |  | *RPS3* | -1049.8 | -1050.3 | 0.5 |
| Epilepsy | Gamma-aminobutyric acid receptor subunit alpha-1 (rs121434579) | *GABBR2* | -1704.2 | -1328.5 | -375.7 |
|  |  | *GABRB2* | -1733.3 | -1639 | -94.3 |
|  |  | *GABRB3* | -1919.4 | -1879.9 | -39.5 |
|  |  | *GABRG2* | -1699.8 | -1707.6 | 7.8 |
|  |  | *HAP1* | -1967.2 | -1630.7 | -336.5 |
|  |  | *PLCL1* | -1481.2 | -1476.3 | -4.9 |
| SZ | D(2) dopamine receptor (rs1801028) | *ADORA2A* | -2139.6 | -2140.8 | 1.2 |
|  |  | *ARRB1* | -1280.2 | -1285.2 | 5 |
|  |  | *ARRB2* | -1828 | -1828.1 | 0.1 |
|  |  | *GNAI1* | -1301.7 | -1301.1 | -0.6 |
|  |  | *GNB1* | -1157.1 | -1157 | -0.1 |
|  |  | *NCS1* | -1333.4 | -2223.5 | 890.1 |

**Figure 3:**
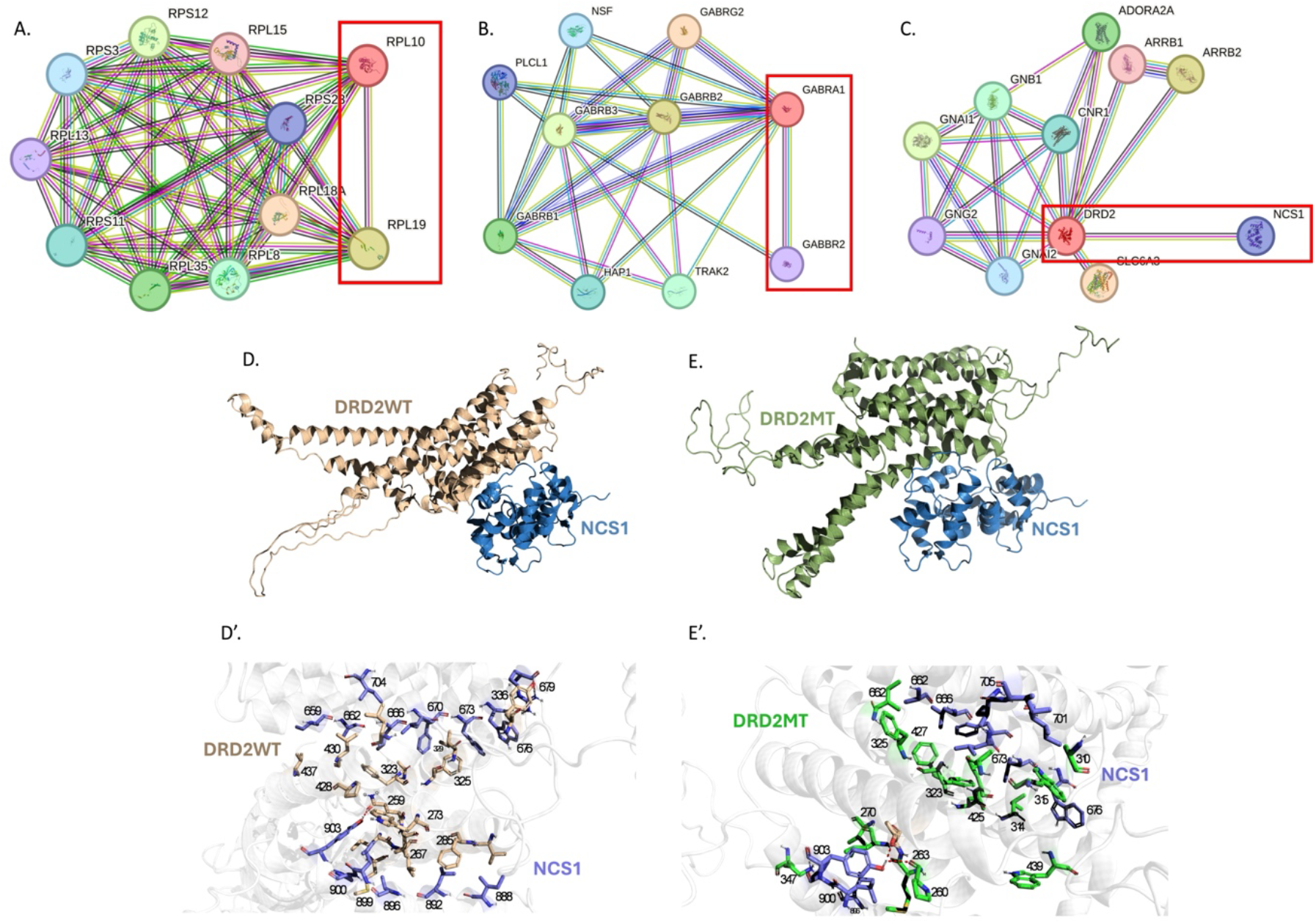
*In silico* prediction of structural changes and protein interaction. Visualization of STRING predicted (confidence level >=0.9%) PPI network of most affected protein of each disorder (Table 1) **(A-C)**. Most affected protein interaction partners are highlighted in red boxes. **(D)** Interaction between WT Dopamine Receptor D2 and Neuronal Calcium Sensor 1, **(D’)** and the interacting residues. **(E, E’)** show the same for MT Dopamine Receptor D2 and Neuronal Calcium Sensor 1. Interacting residues are outlined in **Supplementary Table 2**.

### 3.5 Molecular Docking

It is vital to analyze the change in the protein-protein interaction between the protein of interest and its interacting partner to assess the gravity of the mutation. In the case of DRD2, Ser311 is not involved in the interaction of the protein with NCS1. The mutation causes structural changes dire enough that none of the residues in DRD2 WT model that interact with NCS1, participate in the interaction pair after mutation (Figure 3 D-E’; Supplementary Table 2 A-B). A significant percentage of interaction between WT DRD2 and NCS1 are hydrogen bonds with an average length of 2.44Å. The common type of interaction between MT DRD2 and NCS1 is alkyl/pi-alkyl bond with an average bond length of 4,78Å. Interestingly, there is less formation of salt bridges observed in MT DRD2 and NCS1 than the WT. The introduction of salt bridges into the binding site may have a destabilizing effect on the bonding (30).

Thus, despite not being directly involved in the interaction between DRD2 and NCS1, the Ser311Cys mutation in DRD2 shows a substantiated effect on the interaction between these proteins. This is highlighted by the stark difference in the interacting residues of DRD2 in WT model and MT model.

### 3.6 D(2) dopamine receptor levels and social behavior in SI model of schizophrenia

Post-weaning social isolation in mice (SI mice) results in development of neuropsychiatric like behaviors in adult mice, hence this is considered as valid model to study characteristics of SZ at multiple levels (31). We first assessed the expression levels of D(2) dopamine receptor in hippocampus of SI mice and compared with the group housed (GH) controls. Our analysis reveals a significantly reduced immunofluorescent (IF) intensity in SI mice compared to GH control mice (GH:100%; SI:75%; p= 0.0007; Unpaired *t* test) (Figure 4A, A’, C). Similarly, the expression levels of interaction partner of D(2) dopamine receptor, the NCS1 also reduced in hippocampus of the SI mice (GH: 100%; SI:83%; p= 0.030) (Figure 4B, B’, D).

**Figure 4:**
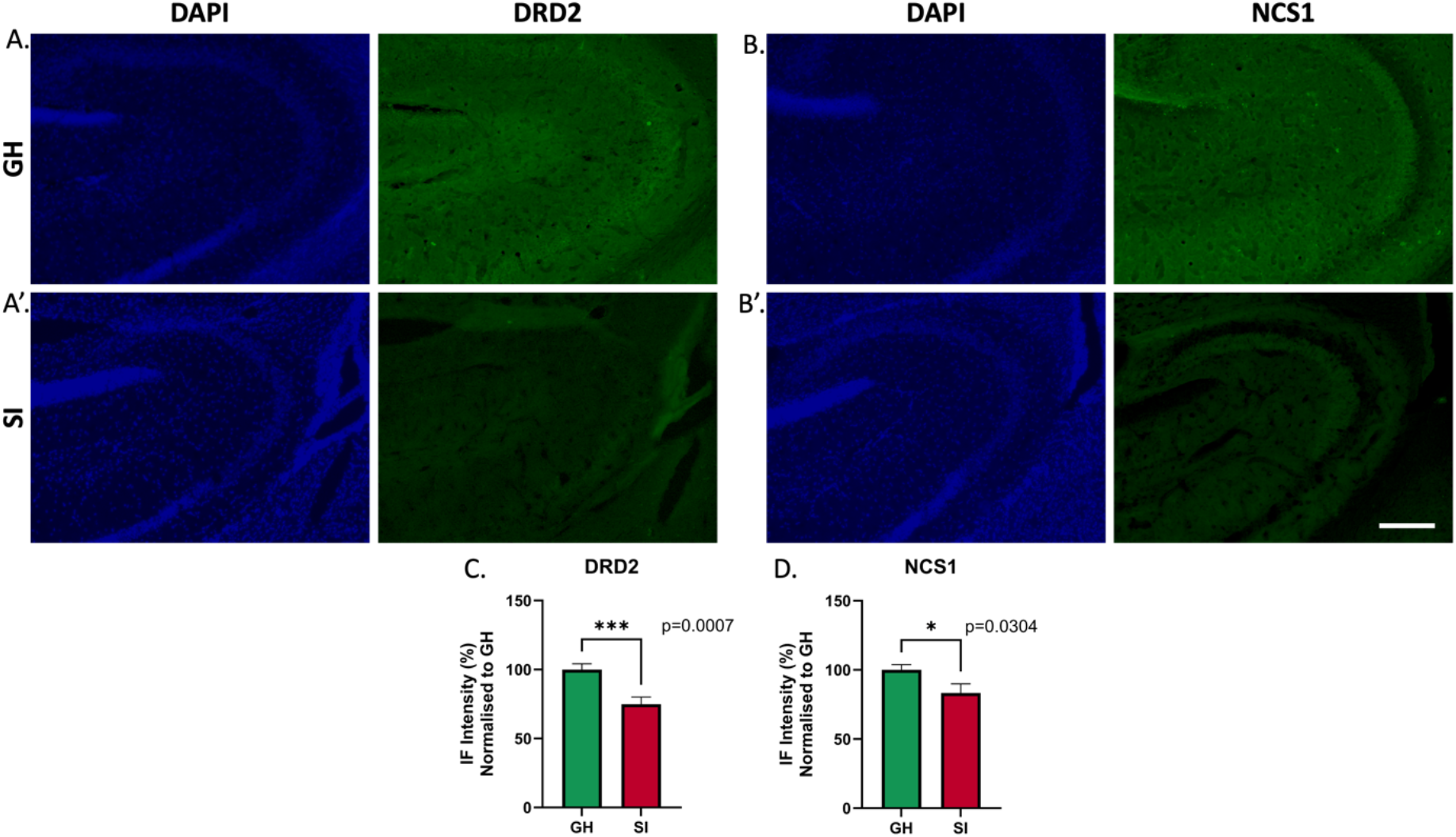
Immunofluorescence analysis of brain sections. Immunoreactivity of DRD2 (Figure **A, A′**), NCS1 (Figure **B, B′**) antibodies in representative hippocampal (CA3 region) sections from group housed (GH) and social isolation (SI) mice. Quantification of immunofluorescent intensity (IF) showing reduction of both DRD2 **(C)** and NCS1 **(D)** in SI compared to GH. Scale bar is 200 μm. Data is represented as Mean±SEM, Students *t*-Test.

Next, we assessed the behavior in these mice in order to find the effect of the reduced expression of D(2) dopamine receptor and NCS1 on social behavior, in particular. In male-female interaction test (Anogenital sniffing; Figure 5A), with (Figure 5Bi: Duration mean; GH: 1.715, SI: 1.125, p=0.2190; student’s *t*-Test; Figure 5Bii: Naïve: GH: 1.980, SI: 1.190, p=0.3216) or without additional stressor (Figure 5Bii: Stressor: GH: 1.449, SI: 1.060, p=0.5390), we observed a trend to towards reduced interaction of SI male mice towards the females in comparison with the GH males, where the reduced interactions occurred during the initial 0-90 seconds, compared to 91-180 seconds (Supplementary Figure 4A, B).

**Figure 5:**
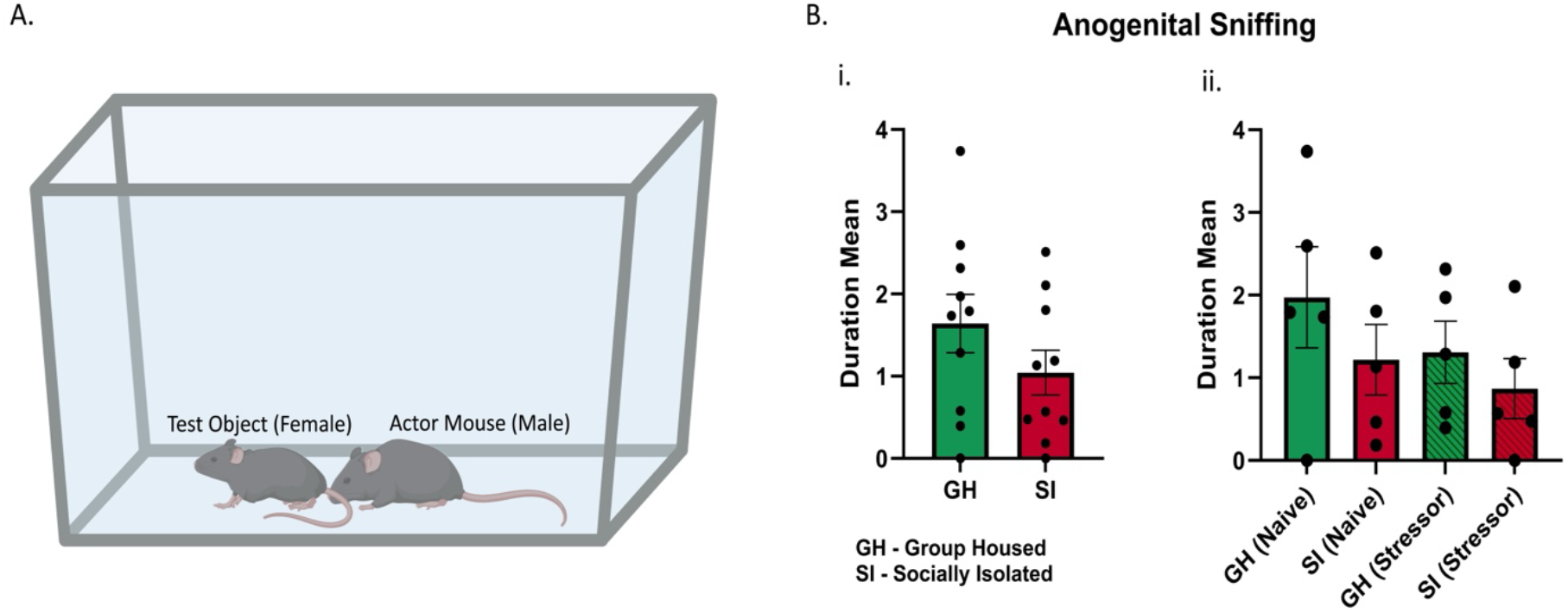
Summary of anogenital sniffing by actor mouse, comparing group housed and social isolated male mice. **(A)** Pictorial representation of anogenital sniffing. **(B)** Summary of duration mean anogenital sniffing performed by actor mouse in total duration of 180 seconds. (i) There is an observed trend of reduction of anogenital sniffing in SI mice in comparison to group housed mice. (ii) To establish the minimized effect of an additional stressor (intraperitoneal administration of 150μl of 1.5% DMSO, striped columns) we also compared these groups segregated. Data is represented as Mean±SEM, Students *t*-Test.

## 4 Discussion

Our *in-silico* analysis demonstrates that known mutations in genes *RPL10, GABRA1* and *DRD2*, associated with ASD, epilepsy and SZ respectively, in Indian population, are the most potentially damaging ones with respect to the protein structure and subsequent protein functions (Figure 1). Synaptic proteins regulate ion channel localization, synaptic vesicle cycle, positional priming and other critical functions (32). Indeed, except for epilepsy conditions, the proteins coded by genes are localized to both pre and post synaptic compartments in ASD and SZ. Of all, the mutations in *DRD2* seems to affect the structural confirmation of the protein, and potentially can cause disturbed synaptic transmission (Figure 2 A-B). Thus, by studying the mutations in these genes, we can understand the mechanism by which these mutations may contribute to the etiology of synaptopathies mentioned above. Further, using *in vivo* model system, we show alterations in D(2) dopamine receptor, its interacting partner NCS1 and a trend towards reduced social interaction (Figure 5B i-ii).

By employing molecular simulation dynamics (MSD), we observed the effect of mutation on interacting amino acid residues of RPL10, GABRA1 and DRD2 with the protein of interest. Thus, based on the MSD analysis, we conclude that mutations in RPL10 (Supplementary Figure 2) and GABRA1 (Supplementary Figure 3) does not drastically impact the structure of the protein, and that the fluctuations are well within normal range. This hints at the fact that despite being in the coding region and a missense SNP, the mutation contributes to the disorder without destabilizing the structure. In the case of DRD2, it can be observed that surprisingly, the MT variant is more rigid than the WT (Figure 2C-RMSD and 2D-RMSF). This is in line with previous study discussing about the SNPs in *DRD2* affect the structural stability, thus hampering the protein-protein interactions (33).

Protein-protein interactions are vital for cellular functions, and disruptions in these interactions can lead to various disorders. Our results reveal, proteins like ion channels and GPCRs can be majorly affected by mutations, and their interactions with other proteins present in their vicinity within the cell or matrix are also impacted (Table 2). Further, our analysis demonstrates the effect of the mutation in *DRD2* on its encoded protein structure and predicted the possibility in alteration of synaptic signaling. This might be possibly mediated by disrupted interaction between DRD2 and NCS1. *FREQ* encodes NCS1 (Neuronal calcium sensor 1 protein), a critical dopamine receptor interacting protein that has a variety of neuronal functions like membrane traffic, cell survival, and ion channel and receptor signaling thus regulating excitability, neuronal development and learning and memory (34–36). Neuronal calcium sensor 1 and D(2) dopamine receptors colocalize at sites of synaptic activity and this interaction has shown to attenuate ligand-induced D(2) dopamine receptor internalization and receptor phosphorylation, enhancing D(2) dopamine receptor signaling (35,37). Altogether our analysis and previous studies predict a strong influence of genetic mutations on protein-protein interactions thus potentially affecting the synaptic functions.

Disrupted functioning or reduced expression of NCS1 along with reduced DRD2 expression shown to affect the dopamine-dependent learning in mice (38,39). Though statistically non-significant, our behavioral data using post-weaning social isolation mouse model of SZ, reveals a trend towards reduced exploration of experimental male mouse towards the female in sociosexual behavior where anogenital sniffing is a crucial phenomenon (40). Although we did not find a significant outcome in social interaction, the *DRD2* have been implicated in sociability or social withdrawal across species (41). In line with previous studies, we identified reduced expression of both NCS1 and DRD2 in hippocampus of our SI model, with a strong reduction in NCS1 in comparison to DRD2 between the SI model and controls. However, postmortem studies on SZ patient brain samples show differential expression pattern of *DRD2* mRNA isoforms between the brain regions implicated in SZ pathology. *D2S* (short isoform) is elevated and *D2L* (long isoform) is reduced in dorsolateral prefrontal cortex (DLPFC), whereas mRNA levels of both these isoforms seems to be affected in opposite direction in hippocampus (42). Similarly, altered expression of NCS1 have been reported in connection with SZ. Elevated expression of NCS1 was identified in DLPFC of postmortem brains of SZ, whereas reduced expression has been reported particularly in lymphocytes of SZ patients (43,44).

In addition to its SZ association, *DRD2* is one of the strongest candidates linked to ASD as well, and targeting its activity alongside whole DRD2 interactome might be a potential therapeutic strategy (45,46). Further studies are required to address the altered interaction between Neuronal calcium sensor 1 and D(2) dopamine receptor, due to the mutation and its relevance to both SZ and ASD and in developing a convergent therapeutic strategies.

## 6 Author Contributions

N.M., P.K.D., S.M., S.K., A.A wrote the initial manuscript. N.M., P.K.D., S.M., S.M., M.R., S.K., N.M., A.A contributed to the data mining and formal analysis. N.M., P.K.D., N.M., K.K., and B.G.A contributed to bioinformatics, simulation analysis and image processing. A.A and B.G.A designed and supervised the study. All the authors contributed to the final version of the manuscript.

## 8 Acknowledgments

Authors acknowledge the support and facilities provided by the Department of Biotechnology, School of Bioengineering at SRM Institute of Science and Technology, Kattankulathur, India.

## 9 Data Availability Statement

The raw data supporting the conclusions of this article will be made available by the authors, upon request.

## References

1. Lepeta K, Lourenco M V., Schweitzer BC, Martino Adami PV., Banerjee P, Catuara-Solarz S, de La Fuente Revenga M, Guillem AM, Haidar M, Ijomone OM, et al. Synaptopathies: synaptic dysfunction in neurological disorders – A review from students to students. J Neurochem (2016) 138:785–805. doi: 10.1111/jnc.13713

2. Hegde PR, Nirisha LP, Basavarajappa C, Suhas S, Kumar CN, Benegal V, Rao GN, Varghese M, Gururaj G. Schizophrenia spectrum disorders in India: A population-based study. Indian J Psychiatry (2023) 65:1223–1229. doi: 10.4103/indianjpsychiatry.indianjpsychiatry_836_23

3. Arora NK, Nair MKC, Gulati S, Deshmukh V, Mohapatra A, Mishra D, Patel V, Pandey RM, Das BC, Divan G, et al. Neurodevelopmental disorders in children aged 2–9 years: Population-based burden estimates across five regions in India. PLoS Med (2018) 15:e1002615. doi: 10.1371/journal.pmed.1002615

4. Singh G, Sharma M, Kumar GA, Rao NG, Prasad K, Mathur P, Pandian JD, Steinmetz JD, Biswas A, Pal PK, et al. The burden of neurological disorders across the states of India: the Global Burden of Disease Study 1990–2019. Lancet Glob Health (2021) 9:e1129–e1144. doi: 10.1016/S2214-109X(21)00164-9

5. Cascella NG, Schretlen DJ, Sawa A. Schizophrenia and epilepsy: Is there a shared susceptibility? Neurosci Res (2009) 63:227–235. doi: 10.1016/j.neures.2009.01.002

6. Jutla A, Foss-Feig J, Veenstra-VanderWeele J. Autism spectrum disorder and schizophrenia: An updated conceptual review. Autism Research (2022) 15:384–412. doi: 10.1002/aur.2659

7. Keller R, Basta R, Salerno L, Elia M. Autism, epilepsy, and synaptopathies: a not rare association. Neurological Sciences (2017) 38:1353–1361. doi: 10.1007/s10072-017-2974-x

8. Fone KCF, Porkess MV. Behavioural and neurochemical effects of post-weaning social isolation in rodents—Relevance to developmental neuropsychiatric disorders. Neurosci Biobehav Rev (2008) 32:1087–1102. doi: 10.1016/j.neubiorev.2008.03.003

9. Li B-J, Liu P, Chu Z, Shang Y, Huan M-X, Dang Y-H, Gao C-G. Social isolation induces schizophrenia-like behavior potentially associated with HINT1, NMDA receptor, and dopamine receptor 2. Neuroreport (2017) 28:462–469. doi: 10.1097/WNR.0000000000000775

10. Koopmans F, van Nierop P, Andres-Alonso M, Byrnes A, Cijsouw T, Coba MP, Cornelisse LN, Farrell RJ, Goldschmidt HL, Howrigan DP, et al. SynGO: An Evidence-Based, Expert-Curated Knowledge Base for the Synapse. Neuron (2019) 103:217-234.e4. doi: 10.1016/j.neuron.2019.05.002

11. Adzhubei I, Jordan DM, Sunyaev SR. Predicting Functional Effect of Human Missense Mutations Using PolyPhen-2. Curr Protoc Hum Genet (2013) 76: doi: 10.1002/0471142905.hg0720s76

12. Schwarz JM, Cooper DN, Schuelke M, Seelow D. MutationTaster2: mutation prediction for the deep-sequencing age. Nat Methods (2014) 11:361–362. doi: 10.1038/nmeth.2890

13. Waterhouse A, Bertoni M, Bienert S, Studer G, Tauriello G, Gumienny R, Heer FT, de Beer TAP, Rempfer C, Bordoli L, et al. SWISS-MODEL: homology modelling of protein structures and complexes. Nucleic Acids Res (2018) 46:W296–W303. doi: 10.1093/nar/gky427

14. Van Der Spoel D, Lindahl E, Hess B, Groenhof G, Mark AE, Berendsen HJC. GROMACS: Fast, flexible, and free. J Comput Chem (2005) 26:1701–1718. doi: 10.1002/jcc.20291

15. Tian C, Kasavajhala K, Belfon KAA, Raguette L, Huang H, Migues AN, Bickel J, Wang Y, Pincay J, Wu Q, et al. ff19SB: Amino-Acid-Specific Protein Backbone Parameters Trained against Quantum Mechanics Energy Surfaces in Solution. J Chem Theory Comput (2020) 16:528–552. doi: 10.1021/acs.jctc.9b00591

16. Bussi G, Donadio D, Parrinello M. Canonical sampling through velocity rescaling. J Chem Phys (2007) 126: doi: 10.1063/1.2408420

17. Martoňák R, Laio A, Parrinello M. Predicting Crystal Structures: The Parrinello-Rahman Method Revisited. Phys Rev Lett (2003) 90:075503. doi: 10.1103/PhysRevLett.90.075503

18. Szklarczyk D, Kirsch R, Koutrouli M, Nastou K, Mehryary F, Hachilif R, Gable AL, Fang T, Doncheva NT, Pyysalo S, et al. The STRING database in 2023: protein–protein association networks and functional enrichment analyses for any sequenced genome of interest. Nucleic Acids Res (2023) 51:D638–D646. doi: 10.1093/nar/gkac1000

19. Kozakov D, Hall DR, Xia B, Porter KA, Padhorny D, Yueh C, Beglov D, Vajda S. The ClusPro web server for protein–protein docking. Nat Protoc (2017) 12:255–278. doi: 10.1038/nprot.2016.169

20. Caroline Gora, Ana Dudas, Lucas Court, Anil Annamneedi, Gaëlle Lefort, Thiago Nakahara, Nicolas Azzopardi, Adrien Acquistapace, Anne-Lyse Laine, Anne-Charlotte Trouillet, et al. Effect of the social environment on olfaction and social skills in WT and a mouse model of autism. Res Sq (2024)

21. Scearce-Levie K, Roberson ED, Gerstein H, Cholfin JA, Mandiyan VS, Shah NM, Rubenstein JLR, Mucke L. Abnormal social behaviors in mice lacking Fgf17. Genes Brain Behav (2008) 7:344–354. doi: 10.1111/j.1601-183X.2007.00357.x

22. Authier N, Dupuis E, Kwasiborski A, Eschalier A, Coudoré F. Behavioural assessment of dimethylsulfoxide neurotoxicity in rats. Toxicol Lett (2002) 132:117–121. doi: 10.1016/S0378-4274(02)00052-8

23. Friard O, Gamba M. <scp>BORIS</scp> : a free, versatile open-source event-logging software for video/audio coding and live observations. Methods Ecol Evol (2016) 7:1325–1330. doi: 10.1111/2041-210X.12584

24. Annamneedi A, del Angel M, Gundelfinger ED, Stork O, Çalışkan G. The Presynaptic Scaffold Protein Bassoon in Forebrain Excitatory Neurons Mediates Hippocampal Circuit Maturation: Potential Involvement of TrkB Signalling. Int J Mol Sci (2021) 22:7944. doi: 10.3390/ijms22157944

25. Baskin B, Gibson WT, Ray PN. Duchenne muscular dystrophy caused by a complex rearrangement between intron 43 of the DMD gene and chromosome 4. Neuromuscular Disorders (2011) 21:178–182. doi: 10.1016/j.nmd.2010.11.008

26. Noble EP. The DRD2 gene in psychiatric and neurological disorders and its phenotypes. Pharmacogenomics (2000) 1:309–333. doi: 10.1517/14622416.1.3.309

27. Maillard P, Baer S, Schaefer É, Desnous B, Villeneuve N, Lépine A, Fabre A, Lacoste C, El Chehadeh S, Piton A, et al. Molecular and clinical descriptions of patients with <scp> GABA _A_</scp> receptor gene variants (<scp>GABRA1</scp>, <scp>GABRB2</scp>, <scp>GABRB3</scp>, <scp>GABRG2</scp>): A cohort study, review of literature, and genotype–phenotype correlation. Epilepsia (2022) 63:2519–2533. doi: 10.1111/epi.17336

28. Klauck SM, Felder B, Kolb-Kokocinski A, Schuster C, Chiocchetti A, Schupp I, Wellenreuther R, Schmötzer G, Poustka F, Breitenbach-Koller L, et al. Mutations in the ribosomal protein gene RPL10 suggest a novel modulating disease mechanism for autism. Mol Psychiatry (2006) 11:1073–1084. doi: 10.1038/sj.mp.4001883

29. Nicholson H, Becktel WJ, Matthews BW. Enhanced protein thermostability from designed mutations that interact with α-helix dipoles. Nature (1988) 336:651–656. doi: 10.1038/336651a0

30. Hendsch ZS, Tidor B. Do salt bridges stabilize proteins? A continuum electrostatic analysis. Protein Science (1994) 3:211–226. doi: 10.1002/pro.5560030206

31. Matsumoto K, Fujiwara H, Araki R, Yabe T. Post-weaning social isolation of mice: A putative animal model of developmental disorders. J Pharmacol Sci (2019) 141:111–118. doi: 10.1016/j.jphs.2019.10.002

32. Gundelfinger ED, Fejtova A. Molecular organization and plasticity of the cytomatrix at the active zone. Curr Opin Neurobiol (2012) 22:423–430. doi: 10.1016/j.conb.2011.10.005

33. Lira SS, Ahammad I. A comprehensive in silico investigation into the nsSNPs of Drd2 gene predicts significant functional consequences in dopamine signaling and pharmacotherapy. Sci Rep (2021) 11:23212. doi: 10.1038/s41598-021-02715-z

34. Mun H-S, Saab BJ, Ng E, McGirr A, Lipina T V., Gondo Y, Georgiou J, Roder JC. Self-directed exploration provides a Ncs1-dependent learning bonus. Sci Rep (2015) 5:17697. doi: 10.1038/srep17697

35. Nakamura TY, Nakao S, Wakabayashi S. Emerging Roles of Neuronal Ca2+ Sensor-1 in Cardiac and Neuronal Tissues: A Mini Review. Front Mol Neurosci (2019) 12: doi: 10.3389/fnmol.2019.00056

36. Fischer TT, Nguyen LD, Ehrlich BE. Neuronal calcium sensor 1 (NCS1) dependent modulation of neuronal morphology and development. The FASEB Journal (2021) 35: doi: 10.1096/fj.202100731R

37. Kabbani N, Negyessy L, Lin R, Goldman-Rakic P, Levenson R. Interaction with Neuronal Calcium Sensor NCS-1 Mediates Desensitization of the D2 Dopamine Receptor. The Journal of Neuroscience (2002) 22:8476–8486. doi: 10.1523/JNEUROSCI.22-19-08476.2002

38. Ng E, Varaschin RK, Su P, Browne CJ, Hermainski J, Le Foll B, Pongs O, Liu F, Trudeau L-E, Roder JC, et al. Neuronal calcium sensor-1 deletion in the mouse decreases motivation and dopamine release in the nucleus accumbens. Behavioural Brain Research (2016) 301:213–225. doi: 10.1016/j.bbr.2015.12.037

39. Ng E, Georgiou J, Avila A, Trought K, Mun H-S, Hodgson M, Servinis P, Roder JC, Collingridge GL, Wong AHC. Mice lacking neuronal calcium sensor-1 show social and cognitive deficits. Behavioural Brain Research (2020) 381:112420. doi: 10.1016/j.bbr.2019.112420

40. Hull EM, Dominguez JM. Sexual behavior in male rodents. Horm Behav (2007) 52:45–55. doi: 10.1016/j.yhbeh.2007.03.030

41. Ike KGO, Lamers SJC, Kaim S, de Boer SF, Buwalda B, Billeter J-C, Kas MJH. The human neuropsychiatric risk gene Drd2 is necessary for social functioning across evolutionary distant species. Mol Psychiatry (2024) 29:518–528. doi: 10.1038/s41380-023-02345-z

42. Kaalund SS, Newburn EN, Ye T, Tao R, Li C, Deep-Soboslay A, Herman MM, Hyde TM, Weinberger DR, Lipska BK, et al. Contrasting changes in DRD1 and DRD2 splice variant expression in schizophrenia and affective disorders, and associations with SNPs in postmortem brain. Mol Psychiatry (2014) 19:1258–1266. doi: 10.1038/mp.2013.165

43. Koh PO, Undie AS, Kabbani N, Levenson R, Goldman-Rakic PS, Lidow MS. Up-regulation of neuronal calcium sensor-1 (NCS-1) in the prefrontal cortex of schizophrenic and bipolar patients. Proceedings of the National Academy of Sciences (2003) 100:313–317. doi: 10.1073/pnas.232693499

44. Torres KCL, Souza BR, Miranda DM, Sampaio AM, Nicolato R, Neves FS, Barros AGA, Dutra WO, Gollob KJ, Correa H, et al. Expression of neuronal calcium sensor-1 (NCS-1) is decreased in leukocytes of schizophrenia and bipolar disorder patients. Prog Neuropsychopharmacol Biol Psychiatry (2009) 33:229–234. doi: 10.1016/j.pnpbp.2008.11.011

45. Annamneedi A, Gora C, Dudas A, Leray X, Bozon V, Crépieux P, Pellissier LP. Towards the convergent therapeutic potential of G protein-coupled receptors in autism spectrum disorders. Br J Pharmacol (2023) doi: 10.1111/bph.16216

46. Chen R, Ferris MJ, Wang S. Dopamine D2 autoreceptor interactome: Targeting the receptor complex as a strategy for treatment of substance use disorder. Pharmacol Ther (2020) 213:107583. doi: 10.1016/j.pharmthera.2020.107583

