## Supplementary material for "Effect of Schizophrenia linked *DRD2* gene mutations and correlation of social behavior with biochemical changes in a post-weaning isolated mouse model": Mukesh and Divi et al., 2024_Supplementary file_bioRxiv-02-11-2024.pdf

### 1.1 Supplementary Figures

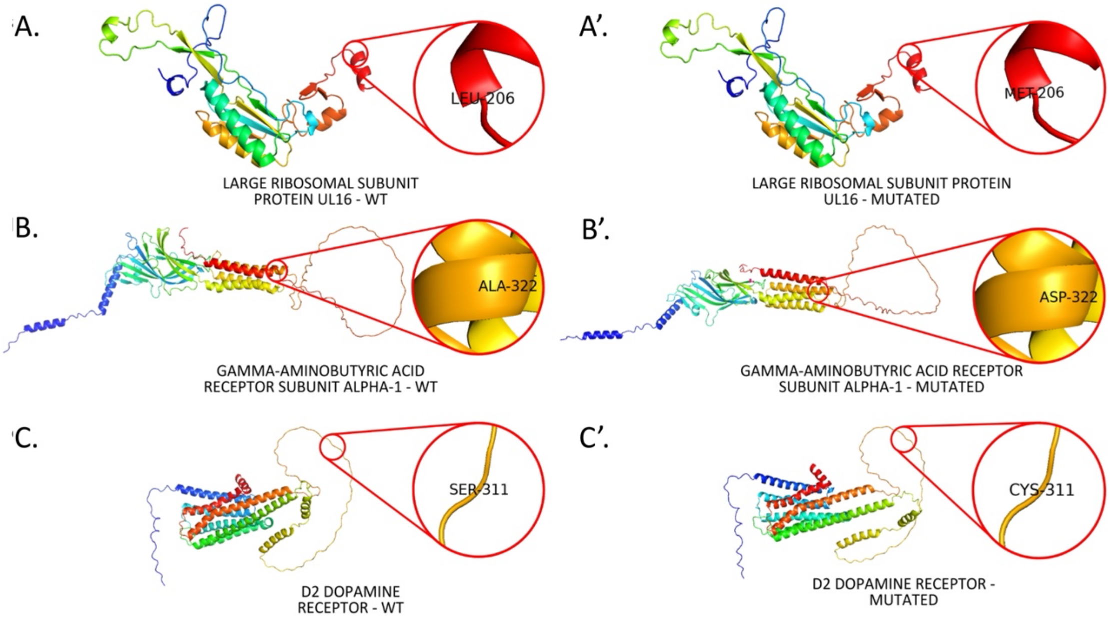


**Supplementary Figure 1: Structural comparison of wild type and mutated proteins.** In RPL10 (A and A'), Leu206 is replaced with Met206. In GABRA1 (B and B') receptor subunit alpha-1, Ala322 is replaced by Asp. In DRD2 (C and C'), Ser311 is substituted by Cys311.

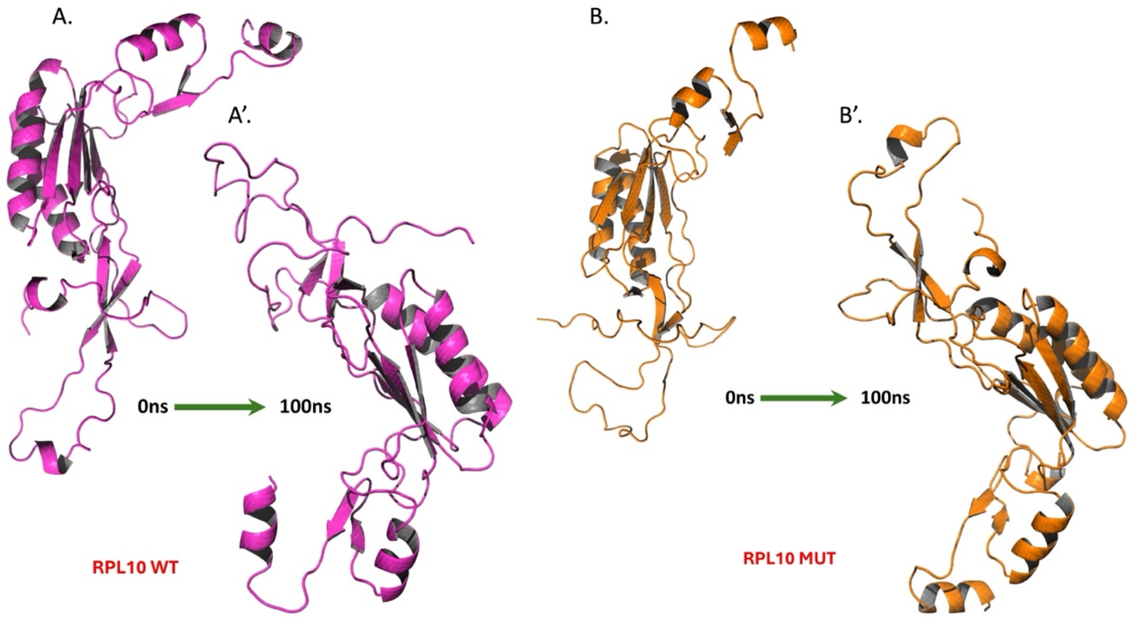


**Supplementary Figure 2: Molecular Simulation Dynamics of RPL10.** The illustration shows the changes in structure between the 0$^{th}$ nanosecond and 100$^{th}$ nanosecond in the simulation of WT (A and A') and MT (B and B') RPL10.

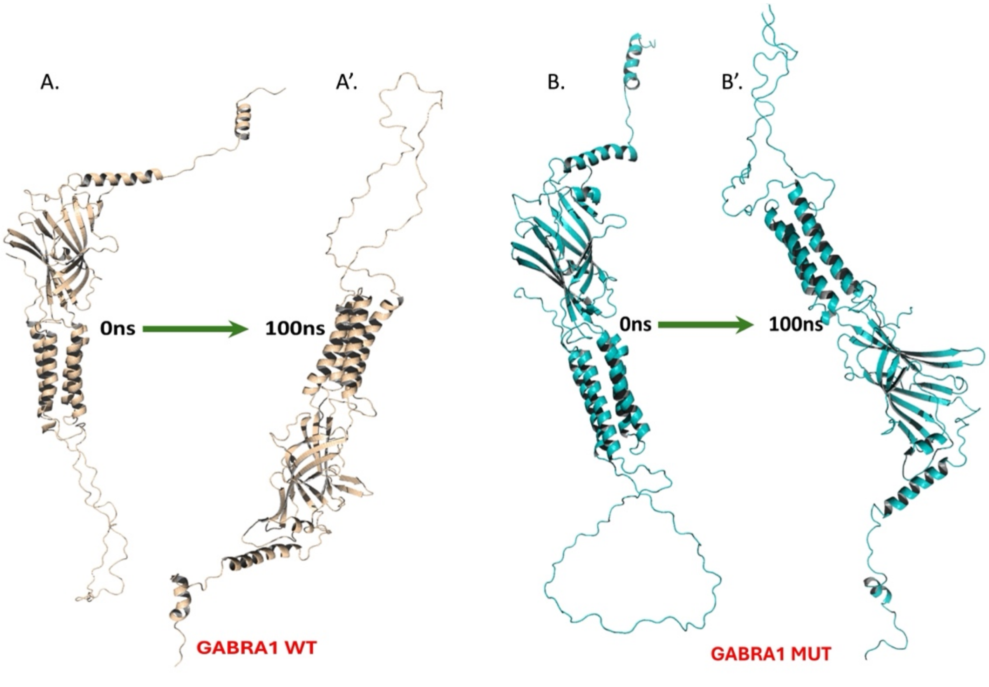


**Supplementary Figure 3: Molecular Simulation Dynamics of GABRA1.** The illustration shows the changes in structure between the 0$^{th}$ nanosecond and 100$^{th}$ nanosecond in the simulation of WT (A and A') and MT (B and B') GABRA1.

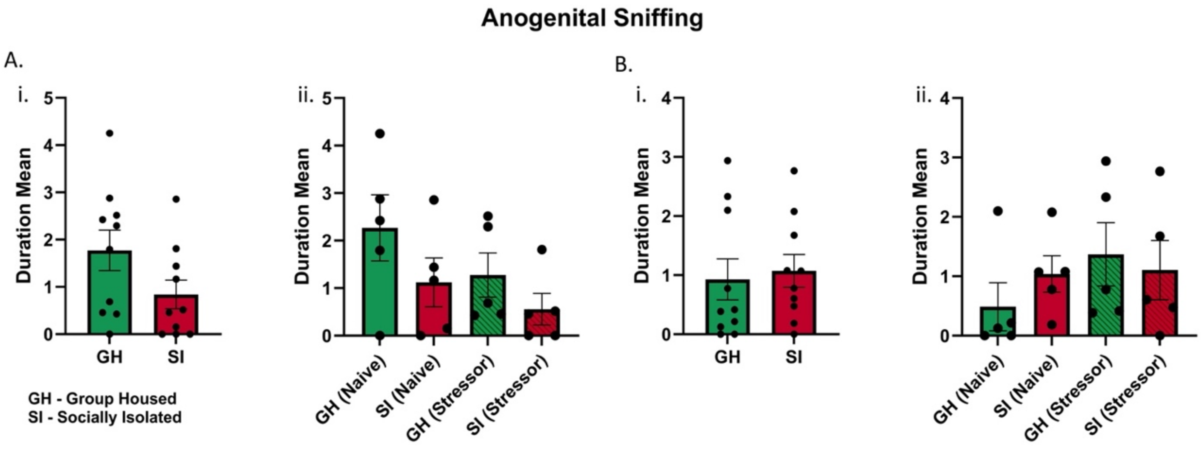


**Supplementary Figure 4: Summary of duration mean anogenital sniffing performed by actor mouse in (A) t=0s → t=90s and (B) t=90s → t=180s. (A-i)** and **(B-i)** summarise the duration mean of anogenital sniffing of group housed and socially isolated mice. **(A-ii)** and further shows the difference between naïve mice and mice subjected to a stressor (intraperitoneal administration of 150ul of 1.5%µl DMSO) **(B-ii)**. Data is represented as Mean±SEM, Students *t*-Test.

### 1.2 Supplementary Tables

**Supplementary Table 1: List of genes collected for this study based on different search engines.** Online sources like PubMed, Google Scholar in Indian population were used to gather data with respect to different neurological conditions.

| DISORDER | CATEGORY | LOCALISATION | INTRONIC | % LOCAL | % TOTAL | EXONIC | % LOCAL | % TOTAL |
|---|---|---|---|---|---|---|---|---|
| Autism Spectrum Disorder | Synaptic | Presynaptic | *HTR2A, SLC6A3* | *33.3%* | *40%* | *RPL10, SLC6A4* | *66.6%* | *60%* |
| | | Postsynaptic | *HRT2A, SLC6A3* | | | *ITGB3, NLGN4Y, RPL10, SLC6A4* | | |
| | Non Synaptic | - | *AVPR1a, DNMT, EN2, TPH2* | *44.5%* | | *MT-ATP6, ND1, ND4, SH3, TPH2* | *55.5%* | |
| Epilepsy | Synaptic | Presynaptic | - | - | *30%* | - | - | *70%* |
| | | Postsynaptic | - | | | *GABRA1, GABRD* | | |
| | Non Synaptic | - | *GST, KCDT7, KCNQ3* | *37.5%* | | *EFHC1, EPM2A, MECP2, NHLRC, RELN* | *62.5%* | |
| Schizophrenia | Synaptic | Presynaptic | *DISC1, DRD2, GRM7, NRG1, SLC6A4* | *85.7%* | *71.4%* | *DRD2* | *14.2* | *28.5%* |
| | | Postsynaptic | *DISC1, DRD2, NRG1,NRGN, SLC6A4* | | | *DRD2* | | |
| | Non Synaptic | - | *CCK-AR, DNMT, NOTCH4, STT3A* | *57.1%* | | *DNMT, GRIK3, XRCC1* | *42.9%* | |

.

**Supplementary Table 2A: Residues involved in PPI of wild-type Dopamine Receptor D2 and wild-type Neuronal Calcium Sensor-1**

| Sl. No. | Interacting Residues Chain A:DRD2 (WT) and Chain B:NCS1 | | Interaction Type | Distance In (Å) |
|---|---|---|---|---|
| 1 | A:GLN54:HE22 | B:TRP160:O | Hydrogen Bond | 2.02948 |
| 2 | A:LYS36:HZ1 | B:THR165:OG1 | Hydrogen Bond | 1.69699 |
| 3 | A:LYS50:HZ1 | B:LEU171:O | Hydrogen Bond | 2.97242 |
| 4 | A:LYS50:HZ3 | B:LEU171:O | Hydrogen Bond | 2.47933 |
| 5 | A:PHE55 | B:ARG294 | Pi-Alkyl | 4.09036 |
| 6 | A:PRO57 | B:LEU82 | Alkyl | 4.81875 |
| 7 | A:TYR186 | B:PRO297 | Pi-Alkyl | 4.53074 |
| 8 | A:TYR186:HH | B:SER296:OG | Hydrogen Bond | 1.83669 |
| 9 | B:ARG292 | A:LEU189 | Alkyl | 5.19277 |
| 10 | B:ARG292:CD | A:THR92:O | Hydrogen Bond | 3.25642 |
| 11 | B:ARG292:HH11 | A:THR92:O | Hydrogen Bond | 1.83207 |
| 12 | B:ARG292:HH12 | A:ASP187:OD1 | Salt Bridge | 1.85001 |
| 13 | B:ARG292:HH12 | A:SER93:O | Hydrogen Bond | 2.75454 |
| 14 | B:ARG292:HH21 | A:GLY188:O | Hydrogen Bond | 1.7646 |
| 15 | B:ARG292:HH22 | A:ASP187:OD1 | Salt Bridge | 1.9118 |
| 16 | B:ARG292:HH22 | A:SER93:O | Hydrogen Bond | 2.94336 |
| 17 | B:ARG292:NH2 | A:ASP187:OD2 | Attractive Charge | 4.90277 |

| | | | | |
|---|---|---|---|---|
| 18 | B:ARG294:HH22 | A:ASP37:OD2 | Salt Bridge | 1.93616 |
| 19 | B:ARG294:NH1 | A:ASP37:OD1 | Attractive Charge | 2.8826 |
| 20 | B:ARG61:CD | A:GLU137:OE1 | Hydrogen Bond | 3.11147 |
| 21 | B:ARG61:HE | A:ASN134:O | Hydrogen Bond | 2.6198 |
| 22 | B:ARG61:HH12 | A:GLU137:OE2 | Salt Bridge; | 2.78926 |
| 23 | B:ARG61:HH21 | A:ASN134:O | Hydrogen Bond | 1.74803 |
| 24 | B:ARG61:HH22 | A:ASN134:OD1 | Hydrogen Bond | 1.95378 |
| 25 | B:ARG61:NH1 | A:GLU137:OE1 | Attractive Charge | 2.88638 |
| 26 | B:ARG61:NH2 | A:GLU137:OE2 | Attractive Charge | 5.24096 |
| 27 | B:CYS168:SG | A:LYS36:O | Hydrogen Bond | 3.66005 |
| 28 | B:HIS303 | A:VAL136 | Pi-Alkyl | 5.37356 |
| 29 | B:LYS439:HZ3 | A:GLU137:OE1 | Salt Bridge | 1.78504 |
| 30 | B:PHE164 | A:ILE51 | Pi-Alkyl | 4.4013 |
| 31 | B:PHE172 | A:PRO39 | Pi-Alkyl | 5.4693 |
| 32 | B:PRO289 | A:LEU189 | Alkyl | 5.18826 |
| 33 | B:PRO297 | A:LEU185 | Alkyl | 4.36294 |
| 34 | B:PRO299 | A:LEU185 | Alkyl | 4.60337 |
| 35 | B:SER60:CB | A:THR135:O | Hydrogen Bond | 3.01778 |
| 36 | B:VAL78 | A:PRO57 | Alkyl | 5.04021 |

**Supplementary Table 2B: Residues involved in PPI of mutant Dopamine Receptor D2 and wild-type Neuronal Calcium Sensor-1**

| Sl. No. | Interacting Residues Chain A:DRD2 (WT) and Chain B:NCS1 | | Interaction Type | Distance In (Å) |
|---|---|---|---|---|
| 1 | A:ARG148:HE | B:ASP400:OD2 | Hydrogen Bond | 1.94059 |
| 2 | A:ARG148:HH22 | B:ASP400:OD1 | Salt Bridge | 1.79729 |
| 3 | A:ARG148:NH1 | B:ASP400:OD2 | Salt Bridge | 4.86869 |
| 4 | A:LEU189:CB | B:TYR192 | Pi-Sigma | 3.76302 |
| 5 | A:PHE55 | B:ILE195 | Pi-Alkyl | 5.49595 |
| 6 | A:PHE56 | B:LEU395 | Pi-Alkyl | 5.48081 |
| 7 | A:PHE58 | B:ILE391 | Pi-Alkyl | 4.54817 |
| 8 | A:PRO57 | B:ILE391 | Alkyl | 4.77905 |
| 9 | A:VAL132 | B:LEU395 | Alkyl | 4.6152 |
| 10 | A:VAL136 | B:ILE403 | Alkyl | 5.0464 |
| 11 | B:ALA188 | A:LEU185 | Alkyl | 3.76986 |
| 12 | B:CYS399 | A:VAL132 | Alkyl | 5.15955 |
| 13 | B:CYS399:SG | A:TYR129 | Pi-Sulfur | 5.6563 |
| 14 | B:CYS401 | A:VAL136 | Alkyl | 3.55933 |
| 15 | B:HIS398 | A:PHE56 | Pi-Pi T-shaped | 5.45605 |
| 16 | B:ILE397 | A:LEU185 | Alkyl | 4.99124 |

| | | | | |
|---|---|---|---|---|
| 17 | B:LEU395 | A:MET131 | Alkyl | 5.23956 |
| 18 | B:TYR192:HH | A:LEU189:O | Hydrogen Bond | 1.91576 |
| 19 | B:TYR199 | A:PHE55 | Pi-Pi Stacked | 4.21955 |
| 20 | B:TYR199:HH | A:GLN54:OE1 | Hydrogen Bond | 1.84439 |
